# Diversity and evolution of the emerging Pandoraviridae family

**DOI:** 10.1101/230904

**Authors:** Matthieu Legendre, Elisabeth Fabre, Olivier Poirot, Sandra Jeudy, Audrey Lartigue, Jean-Marie Alempic, Laure Beucher, Nadège Philippe, Lionel Bertaux, Karine Labadie, Yohann Couté, Chantal Abergel, Jean-Michel Claverie

## Abstract

With DNA genomes up to 2.5 Mb packed in particles of bacterium-like shape and dimension, the first two Acanthamoeba-infecting *Pandoraviruses* remained the most spectacular viruses since their description in 2013. Our isolation of three new strains from distant locations and environments allowed us to perform the first comparative genomics analysis of the emerging worldwide-distributed Pandoraviridae family. Thorough annotation of the genomes combining transcriptomic, proteomic, and bioinformatic analyses, led to the discovery of many non-coding transcripts while significantly reducing the former set of predicted protein-coding genes. We found that the Pandoraviridae exhibit an open pan genome, the enormous size of which is not adequately explained by gene duplications or horizontal transfers. As most of the strain specific genes have no extant homolog and exhibit statistical features comparable to intergenic regions, we suggests that *de novo* gene creation is a strong component in the evolution of the giant Pandoravirus genomes.

## Introduction

For ten years following the serendipitous discovery of the first giant virus (i.e. easily visible by light microscopy) *Acanthamoeba polyphaga* Mimivirus ^1,2^, environmental sampling in search of other Acanthamoeba-infecting viruses only succeeded in the isolation of additional Mimivirus relatives, now forming the still expanding Mimiviridae family ^3–6^. In 2013, we returned to the Chilean coastal area from which we previously isolated *Megavirus chilensis* ^3^, the largest known Mimiviridae. Unexpectedly, we then isolated the even bigger *Pandoravirus salinus* ^7^, the unique characteristics of which suggested the existence of a different family of giant viruses infecting Acanthamoeba. The worldwide distribution of this predicted virus family, the Pandoraviridae, was quickly hinted by our subsequent isolation of *Pandoravirus dulcis*, the first known relative of *P. salinus*, more than 15,000 km away, in a fresh water pond near Melbourne, Australia ^7^. We also spotted Pandoravirus-like particles in an article reporting micrographs of Acanthamoeba infected by an unidentified “endocytobiont” ^8^, the genome sequence of which has recently become available as that of the German isolate *Pandoravirus inopinatum* ^9^.

In the present study, we describe three new members of the Pandoraviridae that were isolated from different environments and distant locations: *Pandoravirus quercus*, isolated from ground soil in Marseille (France); *Pandoravirus neocaledonia*, isolated from the brackish water of a mangrove near Noumea airport (New Caledonia); *Pandoravirus macleodensis*, isolated from a fresh water pond near Melbourne (Australia), only 700 m-away from where we previously isolated *P. dulcis*. In addition to the characterization of their replication cycles in *Acanthamoeba castellanii* by light and electron microscopy, we performed an extensive analysis of the five Pandoravirus strains available in our laboratory through combined genomic, transcriptomic and proteomic approaches (Table 1). We then used these data (together with the genome sequence of *P. inopinatum*) in a comparative manner to build a global picture of the emerging family, as well as to refine the analysis of the gene content of each individual strain. While the number of genes encoding proteins has been revised downwards, we unraveled hundreds of previously unpredicted genes associated to non-coding transcripts. From the comparison of the six representatives at our disposal, the Pandoraviridae family appears quite diverse in terms of gene contents, consistent with an open-ended pan genome. A large fraction of this pan-genome codes for proteins without homologs in cells or other viruses, raising the question of their origin. The purified virions are made of more than 200 different proteins, about half of which are shared by all tested strains in well-correlated relative abundances. This large core proteome is consistent with the highly similar early infection stages exhibited by the different isolates.

**Table 1.**
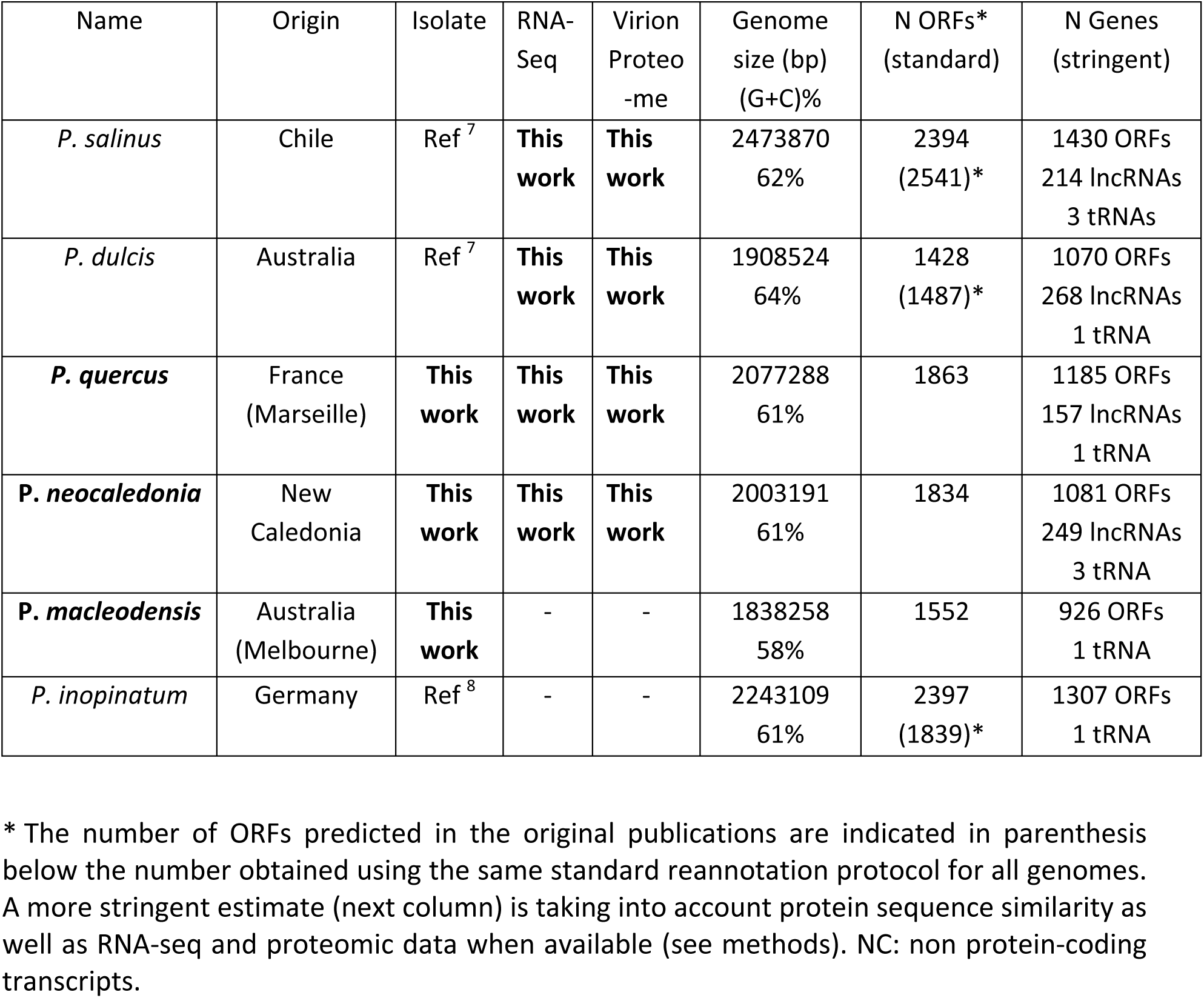
Data on the Pandoravirus isolates used in this work.

| Name | Origin | Isolate | RNA-Seq | Virion Proteome | Genome size (bp) (G+C)% | N ORFs* (standard) | N Genes (stringent) |
| --- | --- | --- | --- | --- | --- | --- | --- |
| *P. salinus* | Chile | Ref <sup>7</sup> | **This work** | **This work** | 2473870<br>62% | 2394<br>(2541)* | 1430 ORFs<br>214 lncRNAs<br>3 tRNAs |
| *P. dulcis* | Australia | Ref <sup>7</sup> | **This work** | **This work** | 1908524<br>64% | 1428<br>(1487)* | 1070 ORFs<br>268 lncRNAs<br>1 tRNA |
| *P. quercus* | France (Marseille) | **This work** | **This work** | **This work** | 2077288<br>61% | 1863 | 1185 ORFs<br>157 lncRNAs<br>1 tRNA |
| *P. neocaledonia* | New Caledonia | **This work** | **This work** | **This work** | 2003191<br>61% | 1834 | 1081 ORFs<br>249 lncRNAs<br>3 tRNA |
| *P. macleodensis* | Australia (Melbourne) | **This work** | - | - | 1838258<br>58% | 1552 | 926 ORFs<br>1 tRNA |
| *P. inopinatum* | Germany | Ref <sup>8</sup> | - | - | 2243109<br>61% | 2397<br>(1839)* | 1307 ORFs<br>1 tRNA |

\* The number of ORFs predicted in the original publications are indicated in parenthesis below the number obtained using the same standard reannotation protocol for all genomes. A more stringent estimate (next column) is taking into account protein sequence similarity as well as RNA-seq and proteomic data when available (see methods). NC: non protein-coding transcripts.

## Results

### Environmental sampling and isolation of new Pandoravirus strains

The sampling and the isolation protocols that led to the discovery of *P. salinus* (from shallow sandy sediment off the coast of central Chile) and *P. dulcis* (from a fresh water pond near Melbourne) have been previously described ^7^. It consists in mixing the sampled material with cultures of Acanthamoeba adapted to antibiotic concentrations high enough to inhibit (or delay) the growth of other environmental microorganisms (especially the fast growing bacteria and fungi). Samples are taken randomly, although from humid environments susceptible to harbor the (rather ubiquitous) Acanthamoeba host. This approach led to the isolation of 3 new Pandoravirus strains described here for the first time: *P. quercus*, *P. neocaledonia*, and *P. macleodensis* (see Methods). Five Pandoravirus strains have thus been isolated and characterized by our laboratory. They exhibit enough divergence to start assessing the conserved features and the variability of the emerging Pandoraviridae family. When appropriate, the present study also includes data from *P. inopinatum*, isolated in a German laboratory from Acanthamoeba infecting a patient with keratitis ^9^.

### Study of the replication cycles and virion ultrastructures

Starting from purified particles inoculated to *A. castellanii* cultures, we analyzed the infectious cycle of each of the new isolates using both light and transmission electron microscopy (ultra-thin section). As previously observed for *P. salinus and P. dulcis*, the replication cycles of these new Pandoraviruses were found to last an average of 12 hours ^7^ (8 hours for the fastest *P. neocaledonia*). The infectious process is the same for all viruses, beginning with the internalization of individual particles by Acanthamoeba cells. Following the opening of the apical pore of the particles, they transfer their translucent content to the cytoplasm through the fusion of the virion internal membrane with that of the phagosome. The early stage of the infection is remarkably similar for all isolates. While we previously reported that the cell nucleus was fully disrupted during the late stage of the infectious cycle ^7^, many more additional observation of all strains revealed neo-synthetized particles in the cytoplasm of cells still exhibiting nucleus-like compartments in which the nucleolus was no longer recognizable (Fig. S1). Eight hours post-infection, mature virions become visible in vacuoles and are released through exocytosis. For all isolates, the replicative cycle ends when the cells lyse and release about a hundred particles, the structures of which do not exhibit any noticeable differences (Fig. 1).

**Figure 1:**
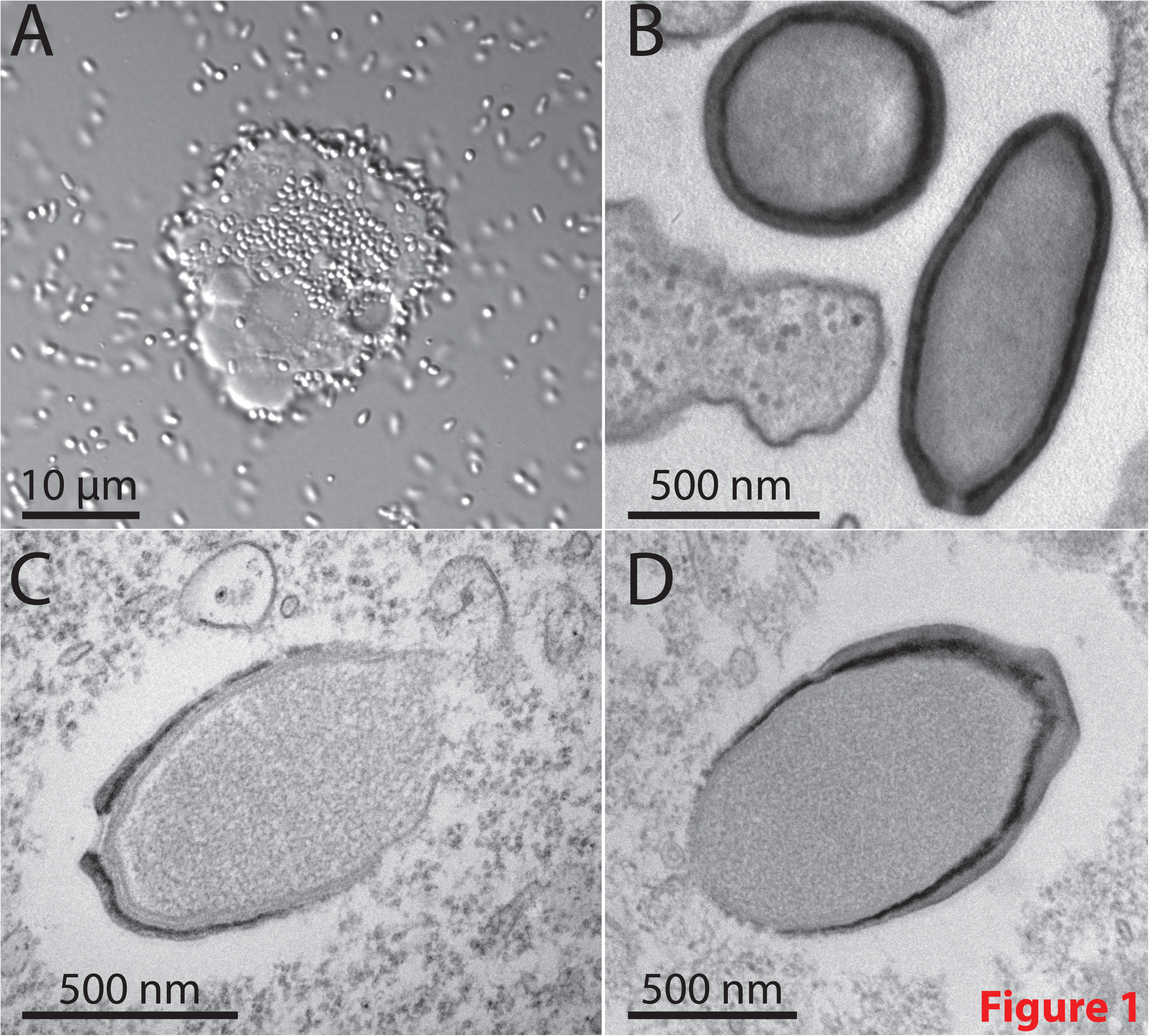
The new Pandoraviruses isolates. (A) Overproduction by an *A. castellanii* cell of *Pandoravirus macleodensis* virions from the environmental sample prior cell lysis. Environmental bacteria can be seen in the culture medium together with *P. macleodensis* virions. (scale bar, 10 µm), (B) TEM image of an ultrathin section of *A. castellanii* cell during the early phase of infection by *P. neocaledonia*. The amoeba pseudopods are ready to engulf the surrounding virions. 10 min pi, virions have been engulfed and are in vacuoles, (C) TEM image of an ultrathin section of *A. castellanii* cell during the assembly process of a *P. salinus* virion and (D) a *P. quercus* virion. Scale bars, 500 nm.

### Genome sequencing and annotation

Genomic DNA of *P. neocaledonia*, *P. macleodensis* and *P. quercus* were prepared from purified particles and sequenced using either the PacBio or Illumina platforms (See Methods). As for *P. salinus*, *P. dulcis* ^7^, and *P. inopinatum* ^9^, the three new genomes assembled as single linear dsDNA molecules (≈60% G+C) with sizes ranging from 1.84 Mb to 2 Mb. In addition to their translucent amphora shaped particles (Fig. 1), higher than average G+C content and genomic gigantism thus remain characteristic features shared by the Pandoraviridae ^7,10^. Given the high proportion of viral genes coding for proteins without database homologs, gene predictions based on purely *ab initio* computational approaches (i.e. “ORFing” and coding propensity estimates) are notoriously unreliable, leading to inconsistencies between teams using different values of arbitrary parameters (e.g. minimal ORF size). Just among families of large dsDNA-viruses infecting eukaryotes, the average protein-coding gene density reportedly varies from one gene every 335 bp (Phycodnaviridae, NCBI: NC_008724) up to one gene every 2120 bp (Herpesviridae, NCBI: NC_003038), while the average consensus is clearly around one gene every kb (such as for bacteria). One thus oscillates between situations where many genes are overpredicted, and others where many real genes are probably overlooked. Such uncertainty about which genes (irrespective of size) are “real” introduces a lot of noise in all subsequent comparative genomic analyses and tests of evolutionary hypotheses. In addition, computational methods are mostly blind to genes expressed as non-protein-coding transcripts.

To overcome the above limitations, we performed strand-specific RNA-seq experiments and particle proteome analyses, the results of which were mapped on the genome sequences. Only genes supported by experimental evidence (or protein similarity) were retained into this stringent reannotation protocol (see Methods, Fig. S2). On one hand, this new procedure led to a reduced set of predicted protein-coding genes, on the other hand it allowed the discovery of an unexpected large numbers of non-coding transcripts (Table 1).

The new set of validated protein-coding genes exhibits a strongly diminished proportion of ORFs shorter than 100 residues, most of which unique to each Pandoravirus strain (Fig. S3). The stringent annotation procedure also resulted in genes exhibiting a well-centered unimodal distribution of Codon Adaptation Index values (Fig. S3).

For consistency, we extrapolated our stringent annotation protocol to *P. inopinatum* and *P. macleodensis*, reducing the number of predicted protein-coding genes taken into account in further comparisons (see Methods, Table 1). The discrepancies between the standard *versus* stringent gene predictions are merely due to the overprediction of small ORFs (length < 300 nucleotides). Such arbitrary ORFs are prone to arise randomly in GC-rich sequences within which stop codons (TAA, TAG, TGA) are statistically less likely to occur by chance than in the non-coding regions of A+T rich genomes. Indeed, the above standard and stringent annotation protocols applied to the A+T rich (74.8%) *Megavirus chilensis* genome ^3^ resulted in two very similar sets of predicted *versus* validated protein-coding genes (1120 *versus* 1108). This control indicates that our stringent annotation protocol is not simply discarding eventually correct gene predictions by arbitrary raising a confidence threshold, but specifically correcting errors induced by the GC-rich composition. Purely computational gene annotation methods are thus markedly less reliable for GC-rich genomes, especially when they encode a large proportion of ORFans, as it is the case for Pandoraviruses. However, it is worth to notice that the fraction of predicted proteins without significant sequence similarity outside of the Pandoravirus family remained quite high (from 67% to 73%, Fig. S4) even after our stringent reannotation.

An additional challenge for the accurate annotation of the Pandoravirus genomes was the presence of introns (virtually undetectable by purely computational methods when they interrupt ORFans). The mapping of the assembled transcript sequences onto the genomes of *P. salinus*, *P. dulcis, P. quercus* and *P. neocaledonia*, allowed the detection of spliceosomal introns in 7.5% to 13% of the validated protein-coding genes. These introns are found in the UTRs as well as in the coding sequences, including on average 14 genes among those encoding the 200 most abundant proteins detected in the particles (see below). These results support our previous suggestion that at least parts of the Pandoravirus transcripts are synthetized and processed by the host nuclear machinery ^7^. Yet, the number of intron per viral gene remains much lower (around 1.2 in average) than for the host genes (6.2 in average, ^11^). Pandoravirus genes also exhibit UTRs in average more than twice as long (Table S1) as those of Mimiviridae ^12^.

In addition, the mapping of the RNA-seq data led to the unexpected discovery of a large number (157 to 268) of long non-coding transcripts (Table 1, Table S1 for detailed statistics). These mRNAs exhibit a polyA tail and are most often transcribed from the reverse strand of validated protein-coding genes, although a smaller fraction are expressed in intergenic (i.e. inter-ORF) regions (Fig. S5). We do not know yet if they play a role in the regulation of the Pandoraviruses genes expression.

About 4% of the non-coding transcripts contain spliceosomal introns. Overall, 82.7% to 87% of the Pandoravirus genomes is transcribed (including ORFs, UTRs and LncRNAs), but only 62% to 68.2% is translated into proteins. Such values are much lower than in giant viruses from other families (e.g. 90% for Mimiviridae ^12^), in part due to the larger UTRs flanking the Pandoravirus genes.

### Comparative genomics

The six protein-coding gene sets obtained from the above stringent annotation procedure were then used as references for different whole genome comparisons aiming to identify specific features of the emerging Pandoraviridae family. Following a sequence similarity-based clustering (see Methods), the relative overlaps of the gene contents of the various strains were computed (Fig. 2A), producing what we will refer now as “protein clusters”.

**Figure 2:**
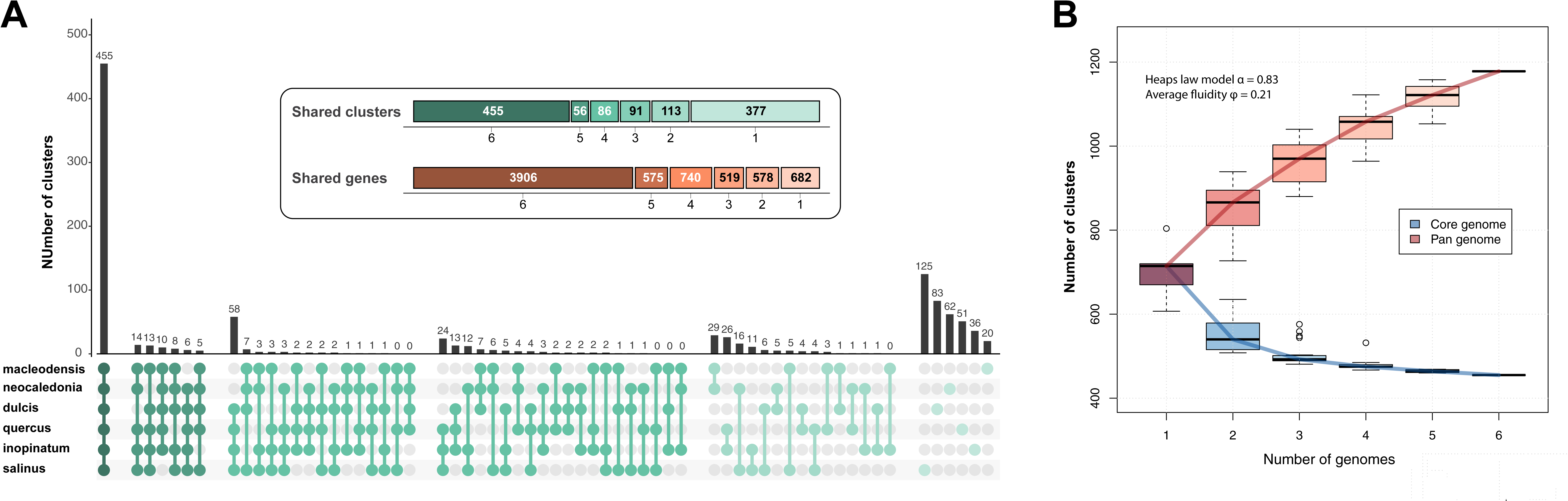
A) Comparison of the Pandoraviruses gene contents. The distribution of all the combinations of shared protein clusters is shown. The inset summarizes the number of clusters and genes shared by 6, 5, 4, 3, 2 and 1 Pandoravirus. B) Core genome and pan genome estimated from the 6 available Pandoraviruses. Estimated heap law α parameter and fluidity are characteristic of open ended pan genomes.

We then computed the number of shared (core) and total genes as we progressively incorporated the genome of the various Pandoravirus isolates into the above analysis, to estimate both the size of the family core gene set, and that of the accessory/flexible gene set. If the 6 available isolates appeared sufficient to delineate a core genome coding for 455 different protein clusters, the “saturation curve” of the total of the different gene families is far from reaching an asymptote, suggesting that the Pandoraviridae pan genome is open-ended, with each additional isolate predicted to contribute more than 50 additional genes (Fig. 2B).

We then investigated the global similarity of the six Pandoravirus isolates by analyzing their shared gene contents both in term of protein sequence similarity and genomic position. The pairwise similarity between the different Pandoravirus isolates ranges from 54% to 88%, as computed from a super alignment of the protein products of the orthologous genes (Table S2). The same data was used to compute a phylogenetic tree that indicates the clustering of the Pandoraviruses into two separate clades (Fig. 3).

**Figure 3:**
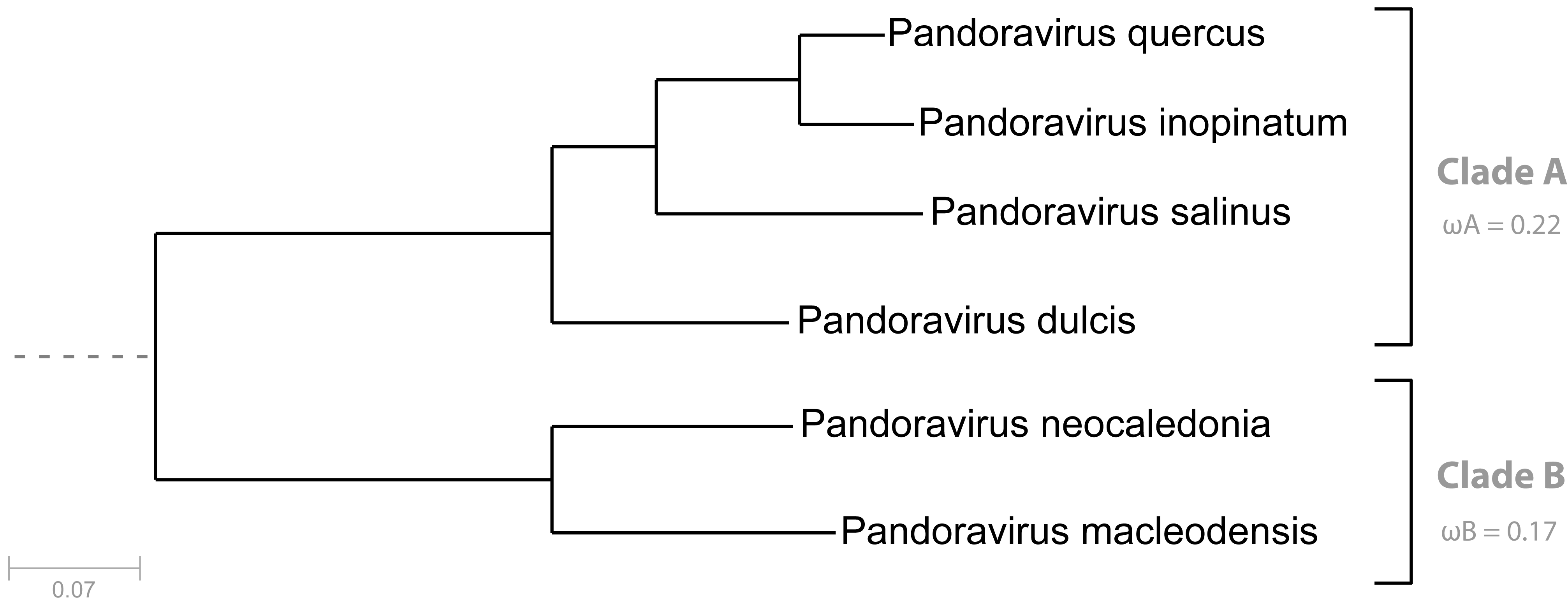
Phylogenetic structure of the Pandoraviridae family. Bootstrap values estimated from resampling are all equal to 1 and so were not reported. Synonymous to non-synonymous substitution rates ratios were calculated for the two separate clades and are significantly different.

Interpreted in a geographical context, this clustering pattern conveys two important properties of the emerging family. On one hand, the most divergent strains are not those isolated from the most distant locations (e.g. the Chilean *P. salinus versus* the French *P. quercus*; the Neo-Caledonian *P. neocaledonia versus* the Australian *P. macleodensis*). On the other hand, two isolates (e.g. *P. dulcis versus P. macleodensis*) from identical environments (two ponds located 700 m apart and connected by a small water flow) were found to be quite different. Pending a larger-scale inventory of the Pandoraviridae, these results already suggest that members of this family are distributed worldwide with similar local and global diversities.

We then analyzed the positions of the homologous genes in the various genomes. We found that despite their divergence at the sequence level (Table S2), 80% of the orthologous genes remain collinear. As shown in Fig. 4, the long-range architecture of the Pandoravirus genomes (i.e. based on the relative positions of orthologous genes) is globally conserved, despite their differences in sizes (1.83 Mb to 2.47 Mb). However, one-half of the Pandoravirus chromosomes (the leftmost region in Fig. 4) curiously appears evolutionary more stable than the other half where most of the non-homologous segments occur. These segments contain strain-specific genes and are enriched in tandem duplications of non-orthologous ankyrin, MORN and F-box motif-containing proteins. Conversely, the stable half of the genome was found to concentrate most of the genes constituting the Pandoraviridae core genome (top of Fig. 4). Interestingly, the local inversion that distinguishes the chromosome of *P. neocaledonia* from the other strains is located near the boundary between the stable and unstable regions, and may be linked to this transition (although it may be coincidental). Finally, all genomes are also enriched in strain specific genes (and/or duplications) at both extremities.

**Figure 4:**
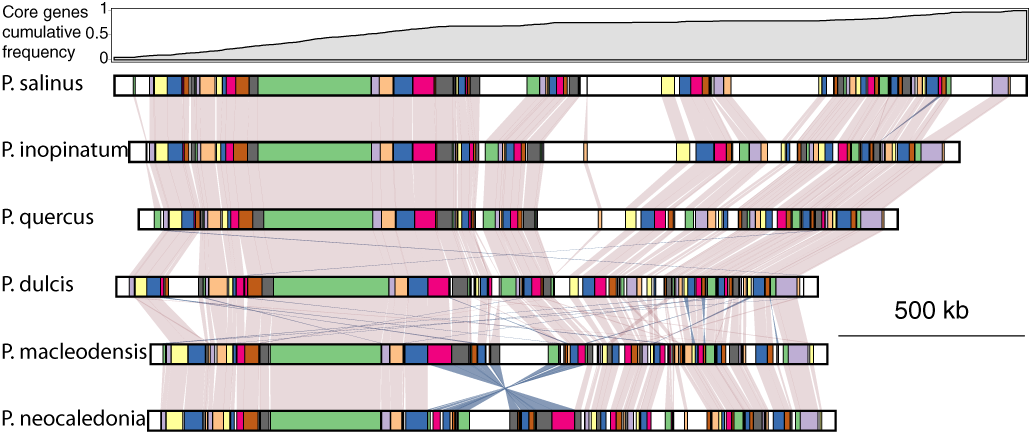
Collinearity of the available Pandoraviruses genomes. Cumulative frequency of core genes is shown at the top. Conserved collinear blocks are colored in the same color in all viruses. White blocks correspond to non-conserved DNA segments.

We then analyzed the distribution of the various Pandoravirus predicted proteins among the standard broad functional categories (Fig. 5). As it is now recurrent for large and giant eukaryotic DNA viruses, the dominant category was by far that of proteins lacking recognizable functional signature. Across the 6 strains, an average of 70% of the protein coding genes corresponded to “unknown function”. Such a high proportion is all the more remarkable as it applies to carefully validated gene sets, from which dubious ORFs have been eliminated. It is thus a biological reality that a large majority of these viral proteins cannot even be vaguely linked to a previously characterized pathway. Remarkably, the proportion of anonymous proteins remains quite high (65%) among the products of the Pandoravirus core genome, that is among the genes, presumably essential, that are shared by the 6 available strains (and probably all future family members, according to Fig. 2B). Interestingly, this proportion remains also very high (≈80 %) among the proteins detected by proteomic analysis as constituting the viral particles. Furthermore, the proportion of anonymous proteins totally dominates the classification of genes unique to each strain, at more than 95 %. The most generic functional category, “protein-protein interaction” is the next largest (from 11.7% to 18.9%), corresponding to the detection of highly frequent and uninformative motifs (e.g. ankyrin repeats). Overall, the proportion of Pandoravirus proteins to which a truly informative function could be attributed is less than 20%, including a complete machinery for DNA replication and transcription.

**Figure 5:**
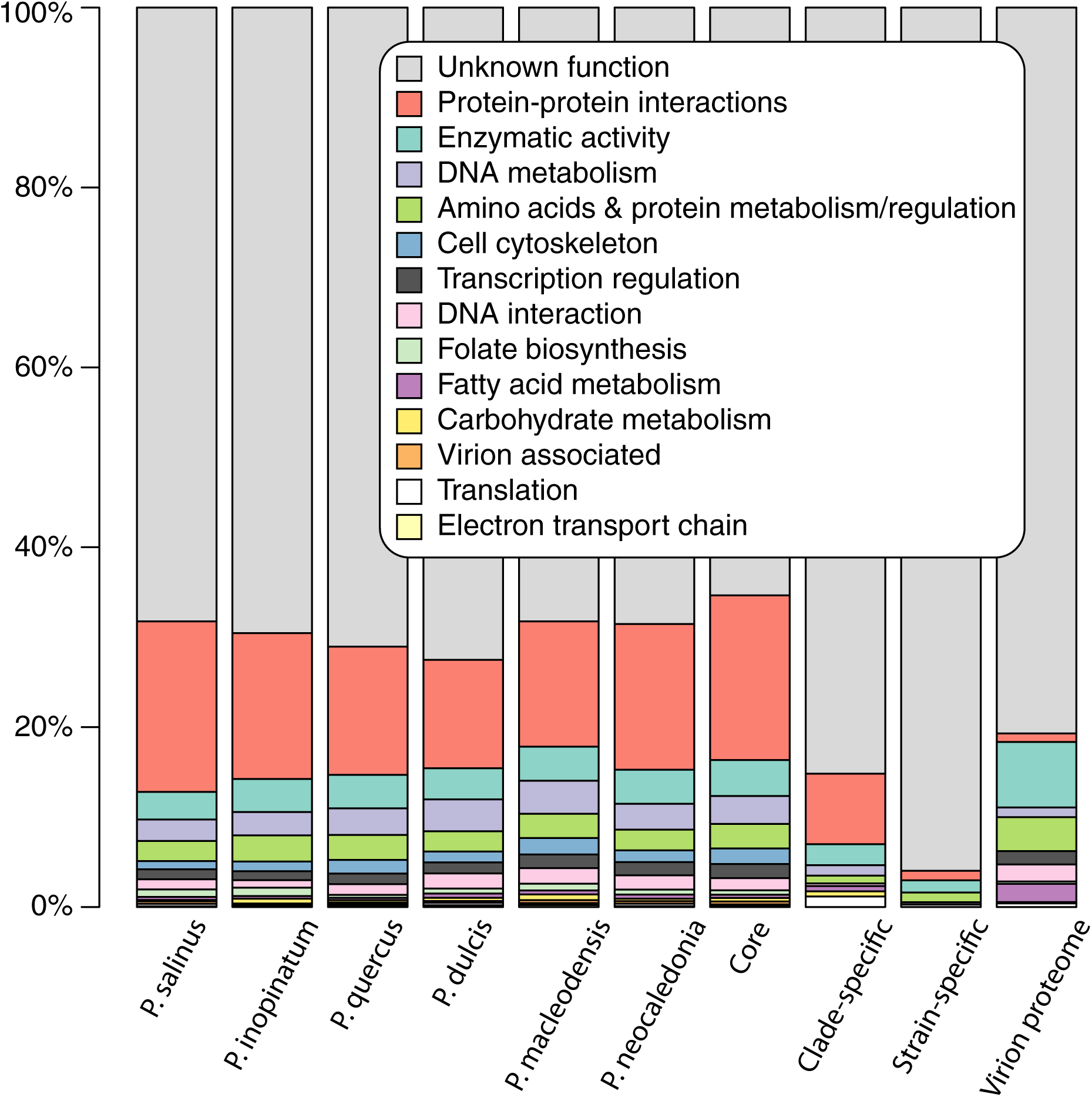
Functional annotations.

We then investigated two evolutionary processes possibly at the origin of the extra-large size of the Pandoravirus genomes: horizontal gene transfers (HGT) and gene duplications. The acquisition of genes by HGT was frequently invoked to explain the extra-large genome size of amoeba-infecting viruses compared to “regular” viruses ^6,13,14^. We computed that up to a third of the Pandoravirus encoded proteins do exhibit sequence similarities (outside of the Pandoravirus family) with proteins from the three cellular domains (Eukarya, Archaea, and Eubacteria) or other viruses (Fig. S4). However, such similarities do not imply that these genes were horizontally acquired. They also could denote a common ancestral origin or a transfer ***from*** a Pandoravirus ***to*** other microorganisms. We individually analyzed the phylogenetic position of each of these cases to infer their likely origin: ancestral – when found outside of clusters of cellular or viral homologs, horizontally acquired – when found deeply embedded in the above clusters, or horizontally transferred to cellular organisms or unrelated viruses in the converse situation (i.e. a cellular protein lying within a Pandoravirus protein cluster). Fig. S6 summarizes the results of this analysis.

We could make an unambiguous HGT diagnosis for 39% of the cases, the rest remaining undecidable or compatible with an ancestral origin. Among the likely HGT, 49% suggested a horizontal gain **by** Pandoraviruses, and 51% the transfer of a gene **from** a Pandoravirus. Interestingly, the acquisition of host genes, a process usually invoked as important in the evolution of viruses, only represent a small proportion (13%) of the diagnosed HGTs, thus less than from the viruses to the host (18%). Combining the above statistics with the proportion of genes (one third) we started from in the whole genome, suggests that at most 15% (and at least 6%) of the Pandoravirus gene content could have been gained from cellular organisms (including 5% to 2% from their contemporary Acanthamoeba host) or other viruses. Such range of values is comparable to what was previously estimated for Mimivirus ^15^. HGT is thus not the distinctive process at the origin of the giant Pandoravirus genomes.

We then investigated the prevalence of duplications among Pandoravirus genes. Fig. 6A compares the proportions of single *versus* duplicated (or more) protein coding genes of the 6 available Pandoraviruses with that computed for representatives of the three other known families of giant DNA viruses infecting Acanthamoeba. It clearly shows that the proportion of multiple-copy genes (ranging from 55% to 44%) is higher in Pandoraviruses, than for the other virus families, although it does not perfectly correlate with their respective genome sizes. The distributions of cluster sizes among the different Pandoravirus strains are similar. Most multiple-copy genes are found in cluster of size 2 (duplication) or 3 (triplication). The number of bigger clusters then regularly decreases with their size (Fig. S7).

**Figure 6:**
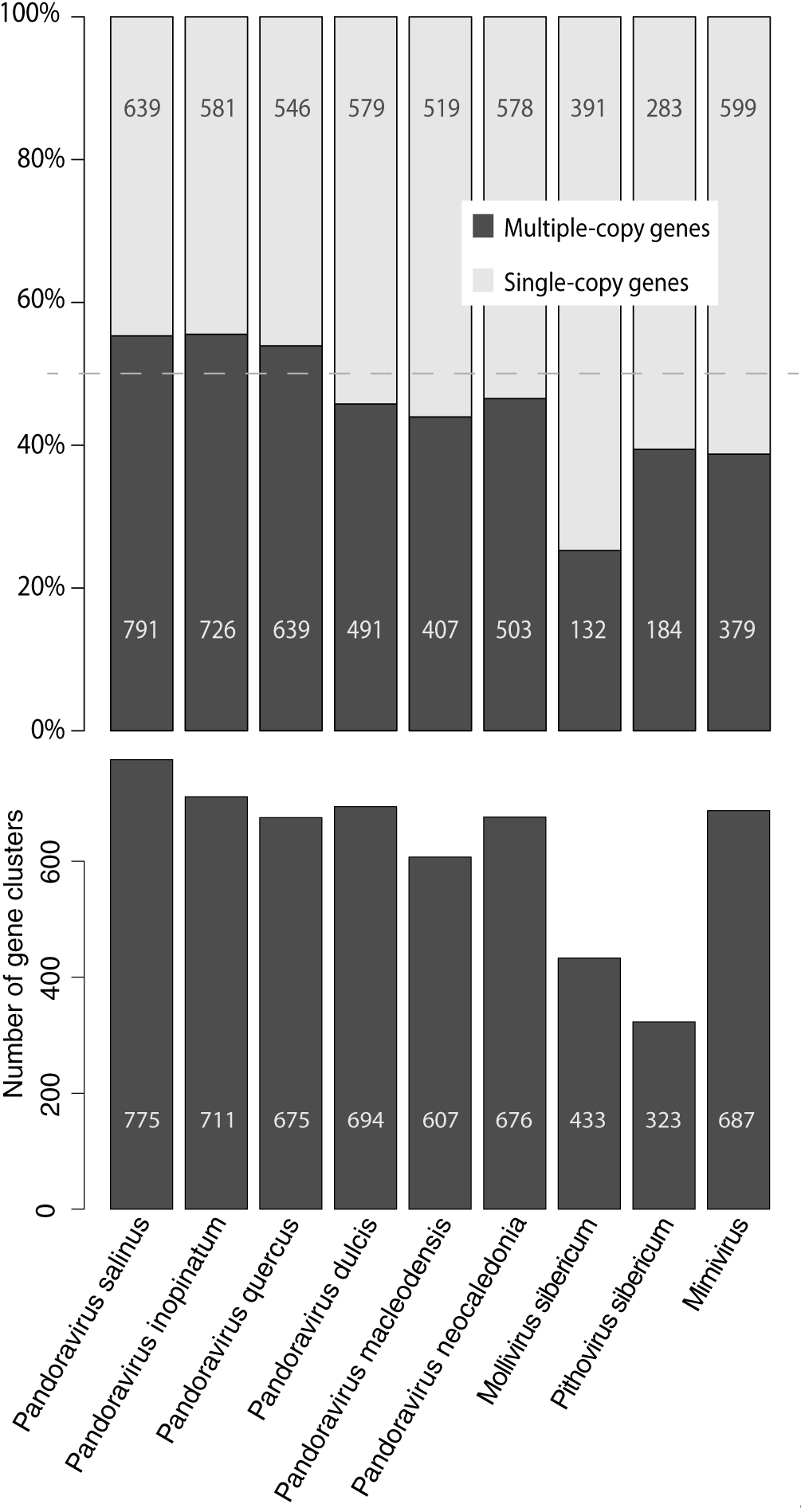
Distribution of single-copy *versus* multiple-copy gene in giant viruses.

Fewer large clusters (size>20) correspond to proteins sharing protein-protein interaction motifs, such as Ankyrin, MORN and F-box repeats. Surprisingly, the absolute number of single copy genes in Pandoraviruses is similar to, and sometimes smaller (e.g. *P. neocaledonia*, 2 Mb) than that in Mimivirus, with a genome (1.18 Mb) half the size. Overall, the number of distinct gene clusters (Fig. 6B) overlaps between the Pandoraviridae (from 607 to 775) and Mimivirus (687), suggesting that, despite their difference in genome and particle sizes, these viruses share comparable genetic complexities.

Gene duplication being such a prominent feature of the Pandoravirus genomes, we investigated it further looking for more insights about its mechanism. First, we computed the genomic distances between every pairs of closest paralogs, most likely resulting from the most recent duplication events. The distributions of these distances, similar for each Pandoravirus, indicated that the closest paralogs are most often located right next to each other (distance =1) or separated by a single gene (distance =2) (Fig. S8).

We then attempted to correlate the physical distance separating duplicated genes with their sequence divergence as a (rough) estimate of their evolutionary distance. We obtained a significant correlation between the estimated “age” of the duplication event and the genomic distance of the two closest paralogs (Fig. S9). These results suggest an evolutionary scenario whereby most duplications are first occurring in tandem, this signal being progressively blurred by subsequent genome alterations events (insertions, inversions, non-local duplications, gene losses).

### Comparative proteomic of Pandoravirus particles (Pandoravirions)

In our initial description of the first Pandoravirus isolates, we presented a mass-spectrometry proteomic analysis of the *P. salinus* particles from which we identified 210 viral gene products, most of which were ORFans or without predictable function. In addition, we detected 56 host (Acantamoeba) proteins. Importantly, none of the components of the virus-encoded transcription apparatus was detected in the particles.

In this work we reproduced the same analyses on *P. salinus*, *P. dulcis*, and two of the new isolates (*P. quercus*, *P. neocaledonia*) to determine to what extent the above features were conserved for members of the Pandoraviridae family with various levels of divergence, and identify the core *versus* the accessory components of a generic Pandoravirion.

Due to the constant sensitivity improvement in mass-spectrometry, our new analyses of purified virions led to the reliable identification of 424 proteins for *P. salinus*, 357 for *P. quercus*, 387 for *P. dulcis*, and 337 for *P. neocaledonia* (see methods). However this increased number of positive identifications corresponds to a range of abundance (intensity-based absolute quantification, iBAQ) values spanning more than five orders of magnitude. Many of the proteins identified in the low abundance tail might thus not correspond to *bona fide* particle components, but to randomly loaded bystanders, “sticky” proteins, or residual contaminants from infected cells. This cautious interpretation is suggested by several observations:

- the low abundance tail is progressively enriched in viral proteins solely identified in the particles of a single Pandoravirus strain (even though several strains possess the homologous genes),
- the proportion of host-encoded proteins putatively associated to the particles also increases at the lowest abundances,
- many of these host proteins were previously detected in purified particles of virus unrelated to the Pandoraviruses but infecting the same host,
- these proteins are abundant in the Acanthamoeba proteome (e.g. actin, peroxidase, etc) and are thus more likely to be retained as purification contaminants.

Unfortunately, the iBAQ value distributions associated to the Pandoravirion proteomes did not exhibit any discontinuity that could serve as an objective abundance threshold to distinguish *bona fide* particle components from dubious ones. However, a plot of the number of identified Acanthamoeba proteins *versus* their rank in the whole proteome showed a rapid increase after rank ≈200 (Fig. S10). Following the same conservative attitude as for the genome reannotation, we decided to disregard the proteins identified after this rank as likely bystanders and only included the 200 most abundant proteins in our further analyses of the particle proteomes (Supplementary dataset S1). Using this stringent proteome definition for each of the four different Pandoravirions, we first investigated the diversity of their constituting proteins and their level of conservation compared to the global gene contents of the corresponding Pandoravirus genomes.

Fig. 7 shows that the particle proteomes include proteins belonging to 194 distinct clusters, 102 of which are shared by the four strains. The core proteome is thus structurally and functionally diverse. It corresponds to 52.6 % of the total protein clusters globally identified in all Pandoravirions. By comparison, the 467 protein clusters encoded by the core genome only represents 41.6 % (i.e. 467/1122) of the overall number of Pandoravirus-encoded protein clusters. The pandoravirus “box” used to propagate the genomes of the different strains is thus significantly more conserved than their gene contents (p<<10^-3^, chi-square test). The genes encoding the core proteome also exhibit the strongest purifying selection among all pandoravirus genes (Fig. S11, D).

**Figure 7:**
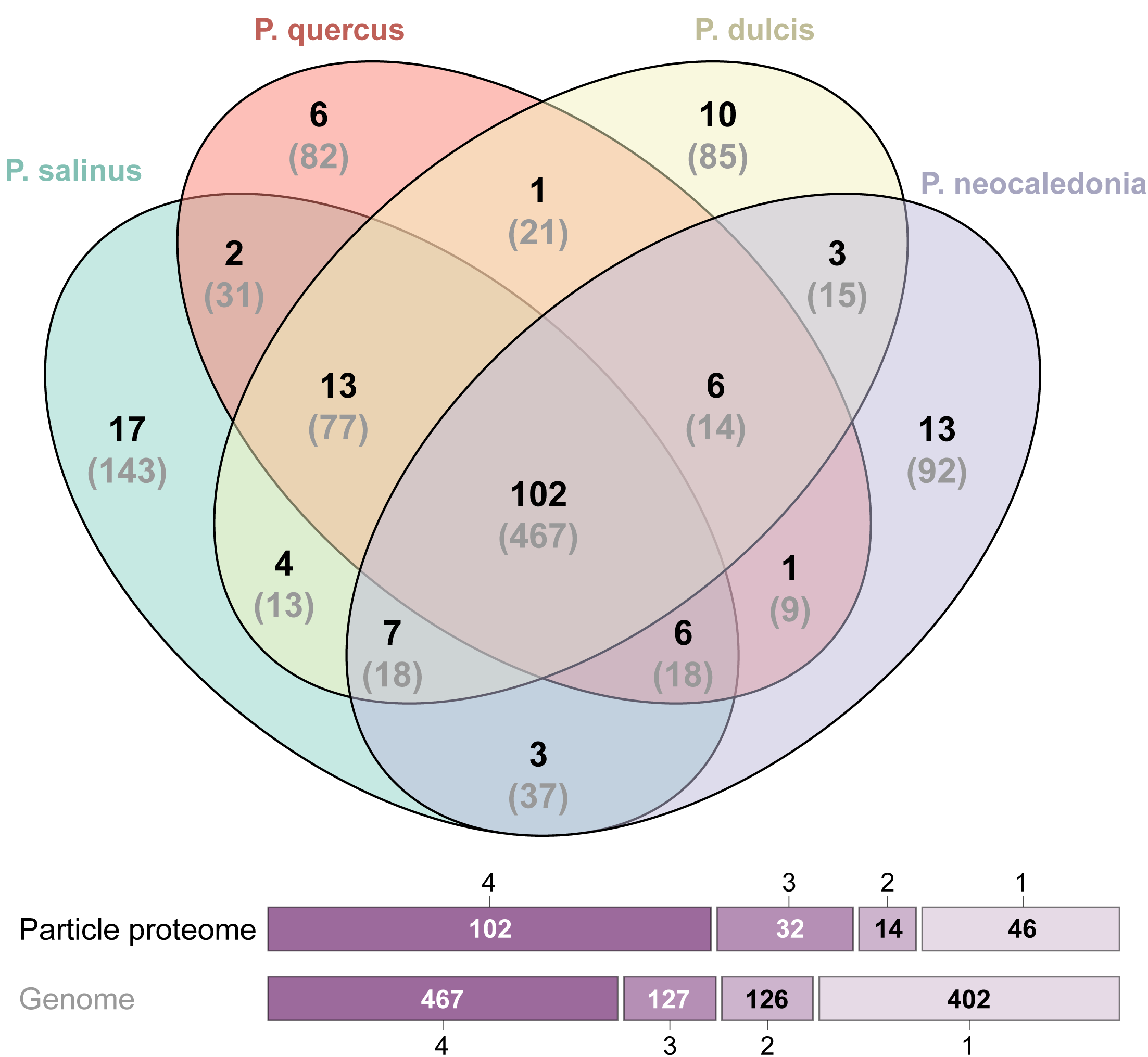
Venn diagram of the particle proteomes of 4 different Pandoravirus strains.

To evaluate the reliability of our proteome analyses we first compared the abundance (iBAQ) values determined for each of the 200 most abundant proteins for two technical replicates and for two biological replicates performed on the same Pandoravirus strain (Fig. S12 A & B). A very good correlation (Pearson’s R > 0.97) was obtained in both cases for abundance values ranging over 3 orders of magnitude. We then compared the iBAQ values obtained for orthologous proteins shared by the virion proteomes of different isolates. Here again, a good correlation was observed (R > 0.81), as expected smaller than for the above replicates (Fig. S12 C & D). These results suggest that although the particles of the different strains appear morphologically identical (Fig. S1), they admit a tangible flexibility both in terms of the protein sets they are made of (with 89% of pairwise orthologues in average), and in their precise stoichiometry.

We then examined the predicted functional attributes of the proteins composing the particles, from the most to the least abundant, hoping to gain some insights about the early infectious process. Unfortunately, only 19 protein clusters could be associated to a functional/structural motif out of the 102 different clusters defining the core particle proteome (Supplementary dataset S2). Such a proportion is less than for the whole genome (Fig. 5), confirming the alien nature of the Pandoravirus particle as already suggested by its unique morphology and assembly process ^7^. These virions are mostly made of proteins without homologs outside of the Pandoraviridae family. No protein even remotely similar to the usually abundant major capsid protein (MCP), a predicted DNA-binding core protein, or a DNA-packaging ATPase, hallmarks of most eukaryotic large DNA viruses, was detected. In particular, a *P. salinus* hypothetical protein (previously ps_862 now reannotated psal_cds_450) recently suggested by Sinclair et al. ^16^ to be a strong major capsid protein candidate was not detected in the *P. salinus* virions, nor its homologs in the other strain proteomes. This result emphasizes the need for the experimental validation of computer predictions made from the “twilight zone” of sequence similarity. No trace of the Pandoravirus-encoded RNA polymerase was detected either, confirming that the initial stage of infection requires the host transcription machinery located in the nucleus. The presence of spliceosomal introns was validated for 56 Pandoravirus genes the products of which were detected in the Pandoravirions (Supplementary dataset S1). This indicates the preservation of a functional spliceosome until the end of the infectious cycle, as expected from the observation of unbroken nuclei (Fig. S1).

Among the 17 non-anonymous protein clusters, four correspond to generic structural motifs and do not bring any functional clue: two collagen-like domains and one Pan/APPLE-like domain that are involved in protein-protein interactions, and one cupin-like domain corresponding to a generic barrel fold. Among the ten most abundant core proteins, nine have no predicted function, except for one exhibiting a C-terminal thioredoxin-like domain (psal_cds_383). It is worth noticing that the predicted membrane-spanning segment of 22 amino acids (85-107) is conserved in all Pandoravirus strains. The 5’UTR of the corresponding genes exhibit 2 introns (in *P. salinus*, *P. dulcis*, and *P. quercus*) and one in *P. neocaledonia*. Thioredoxin catalyzes dithiol-disulfide exchange reactions through the reversible oxidation of its active center. This protein, with another one of the same family (psal_cds_411, predicted as soluble), might be involved in repairing/preventing phagosome-induced oxidative damages to viral proteins prior to the initial stage of infection. The particles also share another abundant redox enzyme, an ERV-like thiol oxidoreductase that may be involved in the maturation of Fe/S proteins. Another core protein (psal_cds_1260), with a remote similarity to a thioredoxin reductase may participate to the regeneration of the oxidized active sites of the above enzymes. Among the most abundant core proteins, psal_cds_232 is predicted as DNA-binding, and may be a major structural component involved in genome packaging. One putative NAD-dependent amine oxidase (psal_cds_628), and one FAD-coupled dehydrogenase (psal_cds_1132) complete the panel of conserved putative redox enzymes. Other predicted core proteins include a Ser/thr kinase and phosphatase that are typical regulatory functions. One serine protease, one lipase, one patatin-like phospholipase, and one remote homolog of a nucleoporin might be part of the toolbox used by the Pandoraviruses to ferry their genomes to the cytoplasm and then to the nucleus (Supplementary dataset S2). Finally, two core proteins (psal_cds_118 and 874) share an endoribonuclease motif and could function as transcriptional regulators targeting cellular mRNA.

At the opposite of defining the set of core proteins shared by all Pandoravirions, we also investigated strain-specific components. Unfortunately, most of the virion proteins unique to a given strain (about 10 in average) are anonymous and in low abundance. No prediction could be made about the functional consequence of their presence in the particles.

## Discussion

We isolated three new strains of Pandoraviruses (*P. neocaledonia*, *P. quercus*, and *P. macleodensis*) from distant locations (resp. New Caledonia, South of France, and Australia) and determined their complete genome sequences. As for the three previously characterized members of this emerging family (*P. salinus* from Chile, *P. dulcis* from Australia and *P. inopinatum* from Germany), their genomes consist in large linear GC-rich dsDNA molecules around 2Mb in size (Table 1). Using the four most divergent pandoravirus strains at our disposal, we combined RNA-seq, virion proteome, and sequence similarity analyses to design and validate a stringent annotation procedure and hopefully eliminate most false positive gene predictions that can both inflate the proportion of ORFans and anonymous proteins, as well as distort the results of comparative analyses. Our - probably over-cautious - gene-calling procedure (reducing the number of predicted protein-coding genes by up to 44%) (Table 1) nevertheless confirmed that an average of 70% of the experimentally validated Pandoravirus genes encode proteins that have no detectable homologue outside of the Pandoraviridae family, and up to 80% for those detected in the particles.

Using the 6 strains known as of today, we also determined that each new member of the family contributed a subset of genes not previously seen in the other genomes, at a rate suggesting that the Pandoraviridae pan genome is open-ended (Fig. 2). Moreover, this flexible (i.e. strain and clade specific) gene content exhibits a much higher proportion of ORFans (respectively 96% and 90%) than the core genome (63%) (Fig. S11, D). According to the usual interpretation, the core genome corresponds to genes that were present in the last common ancestor of a group of viruses while the flexible genome corresponds to genes that appeared since their divergence, through various mechanisms. We then performed further comparative statistical analyses to investigate which mechanisms might be responsible of the large pandoravirus gene content and, possibly, of its continuous expansion.

Based on our conservative predicted gene sets, we determined that gene duplication was a contributing factor in the genome size of Pandoraviruses, with 50% of their genes present in multiple copies (Fig. 6, Fig. S7). However, this value is not vastly different from the proportion (40%) computed for Mimivirus with a genome half the size (Fig. 6). Thus, duplication alone does not explain the much larger gene content (and overall genome size) of the Pandoraviruses. Interestingly, it is neither preferentially expanding the ORfan gene set: the proportion of single-copy ORFans (from 50.7% to 62.7%) compared to those in multiple copies is significantly larger than for non-ORFans (from 30% to 44.5%) (Fischer exact test, p-value <2. 10^-4^). ORFan genes thus tend to be less frequently duplicated.

Horizontal gene transfer (HGT) is also frequently invoked as a mechanism for viral genome inflation ^13,17–20^. Here we estimated that HGT might be responsible for 6 to 15% of the *P. salinus* gene content (Fig. S6). Such a proportion is not exceptional compared to other families of large eukaryotic dsDNA viruses ^15^ with much smaller genomes, and thus does not provide a convincing explanation for the huge gene content of the Pandoraviruses. Furthermore, the large proportion of ORFans among the flexible gene content (Fig. S11) is arguing against their recent acquisition from HGTs, short of postulating that they were taken up from mysterious organisms none of which have yet been characterized and sequenced. Alternatively, one could postulate that the phylogenetic signal from these newly acquired genes might have been erased due to accelerated evolution. However, this is not supported by our data, showing on the contrary that ORFan genes are under strong purifying selection, just to a lesser extent than non-ORFans (Fig. S11). In general, the conceptually appealing notion (inherited from the RNA viruses) that the large proportion of ORFans found in eukaryotic dsDNA viruses would be the consequence of a large mutation rate is progressively losing ground ^21–23^.

To investigate further the origin of the Pandoravirus genes, we performed various statistical analyses in search of what would distinguish genes with orthologs in all strains (core genes) from those only found in strains from the same clade (clade-specific), and from those unique to each strain (strain-specific). To reinforce the assignation of each of the genes to their respective categories, we added a constraint on their genomic positions. For instance, we only considered strain-specific genes found interspersed within otherwise collinear sequences of clade-specific or core genes (Fig S13, A). The genes from the three above categories appeared significantly different with respect to three independent properties (GC content, ORF length and codon adaptation index). More interestingly, the clade-specific and strain-specific genes exhibited average values intermediate between that of the core genes and intergenic sequences (Fig. S13, B-D). The existence of such a gradient unmistakably evocates what is referred to as the *de novo* protein creation (reviewed in ^24–27^). Our data support an evolutionary scenario whereby novel (hence strain-specific) protein-coding genes could randomly emerge from non-coding intergenic regions, then become alike protein-coding genes of older ancestry (i.e. clade-specific and core genes) in response to an adaptive selection pressure (Fig. S11 B). For a long time considered as totally unrealistic on statistical ground ^28^, the notion that new protein-coding genes could emerge *de novo* from non-coding sequences ^29^ started to gain an increasing support following the discovery of many expressed ORFan genes in *Saccharomyces cerevisiae* ^30^, Drosophila ^31^, Arabidopsis ^32^, mammals ^33^, primates ^34^. A different process, called overprinting, involves the use of alternative translation frame from preexisting coding regions. It appears mostly at work in small (mostly RNA-) viruses and bacteria, the dense genomes of which lacks sufficient non-coding regions ^35,36^. However, overprinting would not generate the marked difference in G+C composition between strain-specific and core genes (Fig. S13 B).

There are several good reasons for the eukaryotic-like *de novo* gene creation model to apply to the Pandoraviruses. This evolutionary process requires ORFs that are abundant (to compensate for its contingent nature) and large enough (e.g. > 150 bp) to encode peptides capable of folding into minimal domains (40-50 residues). We previously pointed out that the high G+C content of the Pandoraviruses, compared to the A+T richness of the other known Acanthamoeba-infecting viruses ^10^, statistically increases the size of the random ORFs occurring in non-coding regions. Moreover, these non-coding regions are also much larger in average, representing up to 38% of the total genome (Table S1). The Pandoravirus genomes thus offer an ideal playground for *de novo* gene creation. However, a high G+C genome composition does not imply viral genome inflation and/or an open-ended flexible gene content, as shown by Herpeviruses, another family of dsDNA virus replicating in the nucleus ^37^. Even though HSV-1 and HSV-2 exhibit an overall G+C content of 68% and 70% respectively, their genome remained small (≈150 kb), and their genes coding for core proteins, non-core proteins, as well as their relatively large intergenic regions (≈250±150 bp), do not display any significant difference in composition ^38^. Accordingly, a single gene (US12) has been suggested to have emerged *de novo* ^39^. Thus, Pandoraviruses (and/or their amoebal host) must exhibit some specific features leading them to favor *de novo* gene creation during their evolution. Together, this might be the extensible genome space offered in their particle, the uncondensed state of their DNA genome, the absence of packaged DNA repair enzymes in the virion, or even an unknown template-free machinery generating new DNA. The later mechanism, although highly speculative, appears to be easiest to reconcile with the conserved collinearity of the Pandoravirus genome than intense mutagenesis, duplication, or the shuffling of pre-existing gene. This template-free generation process might be linked to the apparent instability of the right half of the Pandoravirus chromosome, depleted in “core genes” (Fig. 4). We need more genomes to validate the bipartite heterogeneity of the Pandoravirus chromosomes as a distinctive property of the family.

Conceptually, de novo gene creation can occur in two different ways: an intergenic sequence gains transcription before evolving an ORF, or the converse ^26^. The numerous LncRNAs that we detected during the infection cycle of the various Pandoraviruses would appear to bring some support to the transcription first mechanisms. However, most of these non-coding transcripts are antisense of *bona fide* coding regions, and thus could not generate the shift in G+C composition observed for strain-specific genes. Novel proteins could thus emerge from the few intergenic LncRNAs, but mostly from the numerous intergenic (random) ORFs gaining transcription.

The best evidence of *de novo* gene creation, although rarely obtained, is the detection of a significant similarity between the sequence encoding a strain-specific ORFan protein and an intergenic sequence in a closely related strain ^24^. Accordingly, we tested the 318 Pandoravirus strain-specific genes for such occurrences. We found two clearly positive matches. The *P. salinus* psal_cds_1065 (58 aa, 55% GC, CAI=0.287) was found to match a non-coding RNA (pneo_ncRNA_241) in *P. neocaledonia*, and the *P. salinus* psal_cds_415 (96 aa, 54% GC, CAI=0.173) was found to match in an intergenic region in *P. quercus*. In both cases the matches occurred at homologous genomic location. Such low rate of success (yet a positive proof of principle) was expected given the sequence divergence of the available Pandoravirus strains, especially in their intergenic regions.

If we now admit that *de novo* gene creation (as defined above) plays a significant role in the large proportion of strain-specific ORFans and in the open-ended nature of the Pandoraviridae pan genome, it could also have contributed to the pool of family-specific ORFans genes (now shared by 2 to 6 strains) to an unknown extant. The nature of the ancestor of the Pandoraviridae thus remains an unresolved question. Invoking the *de novo* creation hypothesis greatly alleviates the problem encountered when attempting to explain the diversity of the Pandoraviridae gene contents by lineage-specific gene losses and reductive evolution ^10^. Instead of postulating an increasingly complex ancestor as new isolates are exhibiting additional unique genes, we can now attribute them to *de novo* creation. Yet, lineage specific losses can still account for the gene content partially shared among strains.

As seductive as it is, the *de novo* creation hypothesis is nevertheless plagued by its own fundamental difficulties. First, newly expressed (random) proteins have to fold in a compact manner, or at least in a way not interfering with established protein interactions. Although early theoretical studies suggested that stable folding of random amino-acid sequences might be improbable ^40^, several experimental studies have indicated success rate of up to 20% ^41,42^. Thus, globular folding does not appear as a particularly challenging step in the *de novo* creation of non-aggregating proteins. However, high GC content appears to cause encoded proteins to be intrinsically disordered ^43^, thus counter balancing its positive influence on ORF size. Much more problematic is the process by which a random protein would spontaneously acquire a function. For example, only 4 functional (ATP-binding) proteins resulted from the screening of 6x10^12^ random sequences followed by many iterations of in vitro selections and directed evolution ^44^. At the same time, the spontaneous mutation rate of large dsDNA viruses is very low (estimated at less than 10^-7^ substitution per position per infection cycle) ^45^. In absence of a useful function on which to exert a purifying selection, it seems very unlikely that a newly created protein could remain in a genome long enough to acquire a selectable influence on the virus fitness. How the so-called protogene ^30^ manage to be retained during the time they are just a replication burden, remains the dark part of this scenario. Thus, if our comparative genomic studies suggests new hypotheses about the evolution of Pandoraviruses and other giant amoebal viruses, it is far from closing the debate about the genetic complexity of their ancestor ^10,18,20,46,47^.

In the context of this debate, it was previously proposed ^18^ that the Pandoraviruses were highly derived phycodnavirus based on the sole phylogenetic analysis of a handful of genes while disregarding the amazingly unique structural and physiological features displayed by the first two pandoravirus isolates ^7^ as well as the huge number of genes unique to them. Now using the 6 available Pandoravirus genomes, a cladistics clustering based on the presence/absence of homologous genes in the different virus groups robustly separates the Pandoraviridae from the previously established families of large eukaryote-infecting dsDNA viruses (Fig. 8). The only remaining uncertainty concerns the actual position of the yet unclassified Mollivirus sibericum virus that will eventually appear as the seed of a distinct viral family, or as the prototype of smaller pandoravirus relatives from an early diverging branch. More Pandoravidae isolates are needed to delineate the exact boundaries of this new family and resolve the many issues we raised about the origin and mode of evolution of its members.

**Figure 8:**
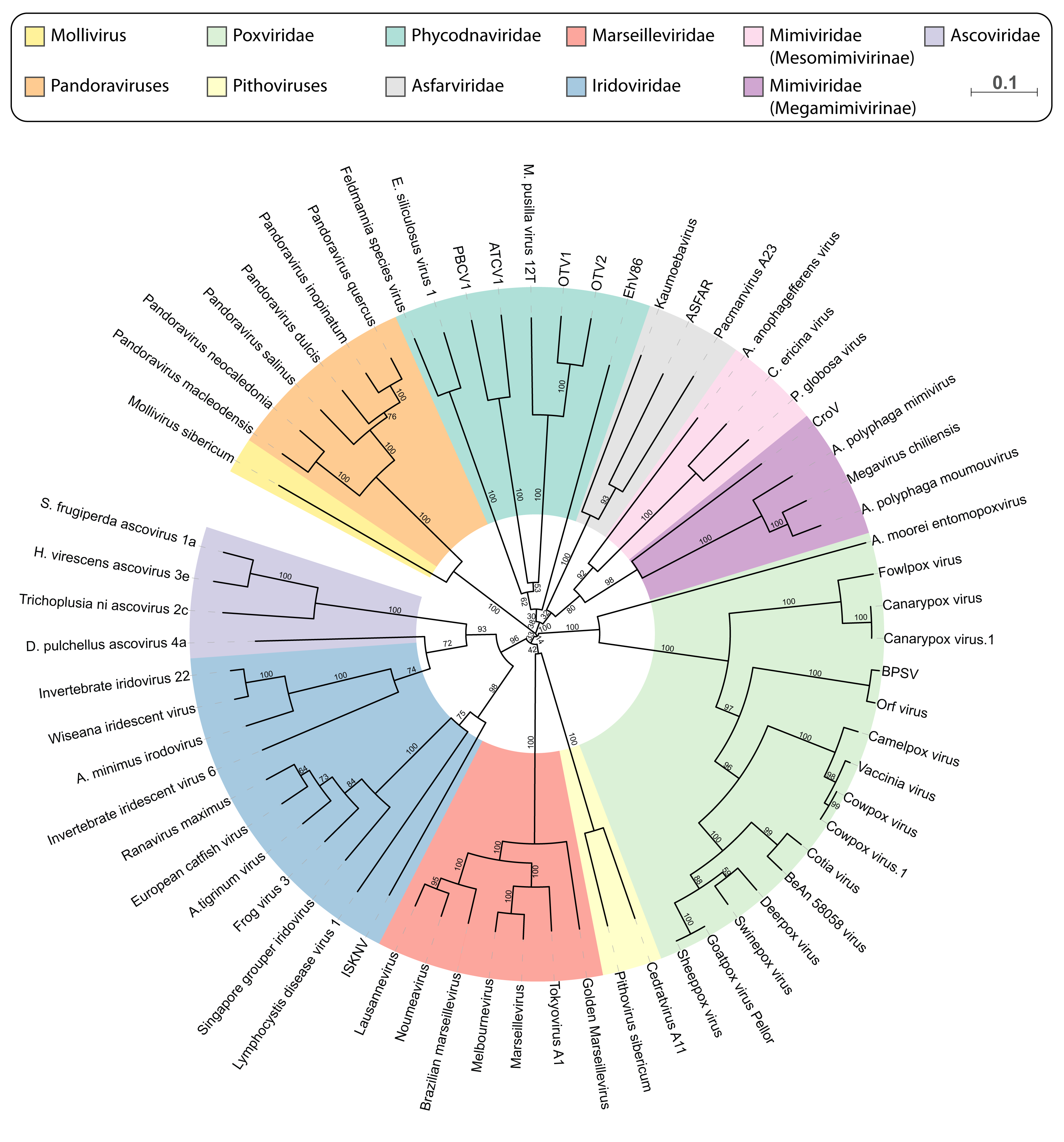
Gene-content based cladistic tree of large DNA viruses. Long virus names have been replaced by acronyms (from top, clockwise). OTV1: *Ostreococcus tauri* virus 1; OTV2: *Ostreococcus tauri* virus 2; EhV86: *Emiliania huxleyi* virus 86; ASFAR: African swine fever virus; CroV: *Cafeteria roenbergensis* virus BV.PW1; BPSV: Bovine papular stomatitis virus; ISKNV: Infectious spleen and kidney necrosis virus.

## Methods

### Environmental sampling and Virus isolation

#### Pandoravirus neocaledonia

A sample from the muddy brackish water of a mangrove near Noumea airport (New-Caledonia, Lat: 22°16'29.50"S, Long: 166°28'11.61"E) was collected. After mixing the mud and the water, 50 ml of the solution was supplemented with 4% of rice media (supernatant obtained after autoclaving 1 L of seawater with 40 grains of rice) and let to incubate in the dark. After one month, 1.5 mL were recovered and 150 μL of pure Fungizone (25 μg/mL final) were added to the sample which was vortexed and incubated overnight at 4°C on a stirring wheel. After decantation, 1 ml supernatant was recovered and centrifuged at 800 × g for 5 min. Acanthamoeba *A. castellanii* (Douglas) Neff (ATCC 30010TM) cells adapted to Fungizone (2.5 μg/mL) were inoculated with 100 μL of the supernatant as previously described ^48^ and monitored for cell death.

#### Pandoravirus macleodensis

A muddy sample was recovered from a pond 700 m away but connected to the La trobe pond in which P. dulcis was isolated ^7^. After mixing the mud and the water, 20 ml of sample were passed through a 20 μm sieve and the filtrate was centrifuged 15 min at 30000 g. The pellet was resuspended in 200 μl of PBS supplemented with antibiotics and 30 μl were added to 6 wells of culture of *A. castellanii* cells adapted to Fungizone (see SI Materials and Methods).

#### Pandoravirus quercus

Soil under decomposing leaves was recovered under an oak tree in Marseille. Few grams were resuspended with 12 mL PBS supplemented with antibiotics. After vortexing 10 min, the tube was incubated during 3 days at 4°C on a stirring wheel. The tube was than centrifuged at 200g 5 minutes and the supernatant was recovered, centrifuged 45 min at 6800g. The pellet was resuspended in 500 μl PBS supplemented with the antibiotics. 50 μl of supernatant and 20 μl of the resuspended pellet were used to infect *A. castellanii* cells adapted to Fungizone. As for P. neocaledonia, and P. macleodensis, visible particles resembling Pandoraviruses were visible in the culture media after cell lysis. All viruses were then cloned using an already described procedure ^49^ prior performing DNA extraction for sequencing and protein extraction for proteomic studies.

Synchronous infections were performed for TEM observation of the infectious cycle. mRNA were extracted from the pooled infected cells prior polyA+ enrichment and sent for library preparation and sequencing to the Genoscope.

### Genome sequencing and assembly

Pandoravirus neocaledonia and Pandoravirus quercus genomes were sequenced using the Pacbio sequencing technology. Pandoravirus macleodensis genome was using the Illumina MiSeq technology with large insert (5-8 kb) mate pair sequences. Details on the strategy used for the genomes assemblies are provided in the SI Materials and Methods.

### Genome sequence stringent annotation

A stringent genome annotation was performed using a combination of *ab initio* gene prediction, strand-specific RNA-seq transcriptomic data, Mass spectroscopy proteomic data as well as protein conservation data. The pipeline used is summarized in Fig. S2 and described in the SI Materials and Methods.

### Proteomic analyses

Virion proteomes were prepared as previously described in ^49^ for mass spectrometry-based label free quantitative proteomics. Briefly, extracted proteins from each preparation were stacked in the top of a 4–12% NuPAGE gel (Invitrogen) before R-250 Coomassie blue staining and in-gel digestion with trypsin (sequencing grade, Promega). Resulting peptides were analysed by online nanoLC-MS/MS (Ultimate 3000 RSLCnano and Q-Exactive Plus, Thermo Scientific) using a 120-min gradient. Three independent preparations from the same clone were analysed for each Pandoravirus to characterize particle composition. Characterization of different clones and technical replicates were performed for *P. dulcis*. Peptides and proteins were identified and quantified as previously described (^49^ and SI Materials and Methods).

### Miscellaneous bioinformatics analyses

A detailed description, of the bioinformatics analyses used for protein clustering, genome rearrangements are detailed in the SI Materials and Methods. Codon Adaptation Index (CAI) was measured using the cai tool from the EMBOSS package ^50^. The reference codon usage was computed from the *Acanthamoeba castellanii* most expressed genes. DNA binding prediction of Pandoraviruses proteins was computed using the DNABIND server ^51^.

### Availability of data and materials

The annotated genomic sequence determined for this work as well as the reannotated genomic sequences have been deposited in the Genbank/EMBL/DDBJ database under the following accession numbers: *P. salinus*: KC977571, *P. dulcis*: KC977570, *P. quercus*: MG011689, *P. neocaledonia* : MG011690, *P. macleodensis*: MG011691. The reannotated *P. inopinatum* genome used in our comparative analyses is provided as supplementary Dataset S1.

The mass spectrometry proteomics data have been deposited to the ProteomeXchange Consortium via the PRIDE ^52^ partner repository with the dataset identifier PXD008167.

All the genomic data, gene annotations and transcriptomic data can be visualized on an interactive genome browser at the following address: http://www.igs.cnrs-mrs.fr/pandoraviruses/

## Acknowledgements

This work was partially supported by the French National Research Agency (ANR-14-CE14-0023-01), France Genomique (ANR-10-INSB-01-01), Institut Français de Bioinformatique (ANR–11–INSB-0013), the Fondation Bettencourt-Schueller (OTP51251), by a DGA-MRIS scholarship and by the Provence-Alpes-Côte-d’Azur région (2010 12125). Proteomic experiments were partly supported by the Proteomics French Infrastructure (ANR-10-INBS-08-01) and Labex GRAL (ANR-10-LABX-49-01). We thank the support of the discovery platform and informatics group at EDyP.

## Authors’ contributions

M.L., Y.C., C.A. and J.-M.C. conceived and designed the research; E.F., S.J., A.L., J.-M. A., L. Beucher, N.P., L. Bertaux, Y.C. performed experimental research; M.L, O.P., C.A. and J.-M. C. performed bioinformatics research; L. Beucher, K.L. and Y.C. contributed new reagents/analytic tools; M.L., C.A. and J.-M. C. analyzed data; M.L., C.A. and J.-M. C wrote the paper.

